# Localization and phosphorylation in the Snf1 network is controlled by two independent pathways

**DOI:** 10.1101/2021.06.14.448401

**Authors:** Linnea Österberg, Niek Welkenhuysen, Sebastian Persson, Stefan Hohmann, Marija Cvijovic

## Abstract

AMPK/SNF1 is the master regulator of energy homeostasis in eukaryotic cells and has a key role in glucose de-repression. If glucose becomes depleted, Snf1 is phosphorylated and activated. Activation of Snf1 is required but is not sufficient for mediating glucose de-repression indicating a second glucose-regulated step that adjusts the Snf1 pathway. To elucidate this regulation, we further explore the spatial dynamics of Snf1 and Mig1 and how they are controlled by concentrations of hexose sugars. We utilize fluorescence recovery after photobleaching (FRAP) to study the movement of Snf1 and how it responds to external glucose concentrations. We show that the Snf1 pathway reacts to the presence of glucose. Furthermore, we identify a negative feedback loop regulating Snf1 activity. Our data offer insight into the true complexity of regulation of this central signaling pathway by one signal (glucose depletion) interpreted by the cell in different ways.

## Introduction

AMPK and its yeast homolog SNF1 is the master regulator of energy homeostasis in eukaryotic cells (Hardie 2014; Hardie, Ross, and Hawley 2012). The AMPK/SNF1 family of protein kinases is regulated by multiple stimuli that signal an energy depletion or a significant rise in energy demand. In the yeast *Saccharomyces cerevisiae* the primary function of SNF1 is adaptation to glucose limitation when the use of alternative carbons sources is needed to achieve growth and proliferation (Hedbacker and Carlson 2008). In addition, a broad spectrum of downstream effects, such as lipid biogenesis and gluconeogenesis, is affected by the SNF1 pathway to balance the energy demand and supply (Usaite et al. 2009; J. Zhang, Olsson, and Nielsen 2010).

The Snf1 kinase is constitutively activated by three upstream kinases Elm1, Sak1 and Tos3 (García-Salcedo et al. 2014; Hong et al. 2003; Nath, McCartney, and Schmidt 2003). When a high energy-yield sugar becomes available, such as the hexose sugars glucose, fructose or mannose, Snf1 is rapidly dephosphorylated by the PP1 phosphatase Reg1/2-Glc7, Sit4 or Ptc2 (Amparo Ruiz, Xu, and Carlson 2013; A. Ruiz, Xu, and Carlson 2011; Y. Zhang et al. 2011). The catalytic unit Snf1 alone is not sufficient to mediate glucose de-repression. For stable Snf1 activity, two more proteins need to bind Snf1 to form the heterotrimeric kinase complex SNF1 (Celenza, Eng, and Carlson 1989; M. C. Schmidt and McCartney 2000). The SNF1 complex consists of the catalytic alpha-subunit Snf1, regulatory gamma-subunit Snf4 and a beta-subunit, which can either be Gal83, Sip1 or Sip2 (Jiang and Carlson 1997; M. C. Schmidt and McCartney 2000). The binding of ADP to Snf4 protects from dephosphorylation and inactivation of Snf1 (Chandrashekarappa, McCartney, and Schmidt 2013; F. V. Mayer et al. 2011). ATP competes with ADP for these binding sites, and this competition functions as an energy sensor (F. V. Mayer et al. 2011). The subcellular localization of the complex is regulated by the beta-subunits (Vincent et al. 2001). Localization studies of the three isoforms under high glucose conditions showed that all the beta-subunits seem to reside in the cytosol. With ethanol as the sole energy source, the Sip1 isoform is associated with the vacuolar membrane. Sip2 is located in the cytoplasm, and Gal83 accumulates in the nucleus (Chandrashekarappa et al. 2016; Vincent et al. 2001). Under the shift from high glucose concentrations to ethanol as the sole carbon source, a major proportion of Snf1 and Snf4 localizes together with Gal83 to the nucleus (Vincent et al. 2001). To alter gene transcription in the cell, Snf1 phosphorylates several transcription factors, among which Mig1 is the most prominent (Ostling and Ronne 1998; Michelle A. Treitel, Kuchin, and Carlson 1998). Mig1 in the unphosphorylated state locates to the nucleus and interacts with Cyc8/Ssn6 and Tup1 to repress transcription of glucose repressed genes (Keleher et al. 1992; M. A. Treitel and Carlson 1995). When the primary energy sources are depleted, Snf1 phosphorylates Mig1 on at least four sites (DeVit and Johnston 1999; Michelle A. Treitel, Kuchin, and Carlson 1998). The localization of the SNF1 isoforms have been shown necessary in response to alkaline stress, but all isoforms can phosphorylate Mig1 in response to glucose depletion (Chandrashekarappa et al. 2016). While some roles of the beta-subunits seem to be redundant, especially for Gal83 and Sip2, the subunits cannot complement each other completely (J. Zhang, Olsson, and Nielsen 2010). A proposed model, where the Gal83 isoform of SNF1 has a structural role in the repression complex of SUC2, a gene co-regulated by Mig1 and Mig2 has been formulated (Vega et al. 2016). Still the importance of the SNF1 location driven by different isoforms remains unclear.

To further characterize the role of the different SNF1 isoforms, we utilize FRAP to study the nuclear-cytoplasmic shuttling of Snf1 and how it responds to external glucose concentrations. We show that the Snf1 pathway reacts to the presence of glucose. We identify a negative feedback loop regulating Snf1 activity, as well as define distinct kinetic behavior in Snf1 nucleocytoplasmic shuttling.

## Results

### The kinetics of Snf1 nucleocytoplasmic shuttling is driven by carbon source availability

It is unclear how the Snf1 dynamic spatial distribution contributes to Snf1’s role in the glucose derepressing pathway. To further understand how Snf1 mechanistically regulates energy balance in the cell, we employed fluorescence recovery after photobleaching (FRAP). Exponentially grown yeast cells, grown in YNB with 2 % glucose, with a Snf1-GFP construct were exposed to YNB under three different conditions: 2% glucose, 0.05% glucose or 2% glycerol for at least 1.5 h before the onset of the experiment. The fluorescent Snf1 in the nucleus is bleached, and the subsequent recovery of fluorescence in the nucleus is observed. The FRAP data were analyzed with a non-linear mixed effect framework (NLME), assuming both a single (Figure 1) and a double (SI data files 2, 3 and 4) exponential model. Non-linear mixed-effects modelling is typically used for longitudinal data exhibiting both within and between-subject variability (Davidian and Giltinan 2003). This method has been widely used in pharmacokinetics and pharmacodynamic studies (Lavielle and Mentré 2007; Sissoko et al. 2016), but in recent years it is exploited in single-cell time-lapse data facilitating our understanding of cell-to-cell variability (Almquist et al. 2015; Llamosi et al. 2016; Persson et al. 2020; Welkenhuysen et al. 2017). When analyzing fluorescence measurements of a tagged protein in single cells over time, the observed intensity will differ between measurements even if the cells are in a steady-state due to the measurement error. Moreover, owing to extrinsic variability, cells will have different intensity levels. Using a mixed-effects framework, the observed cell-to-cell variability can be accounted for in the analysis by letting the rate parameters vary between cells according to a probability distribution. Furthermore, a mixed-effects framework allows the assessment of potential correlations between parameters in different cells.

**Figure 1:**
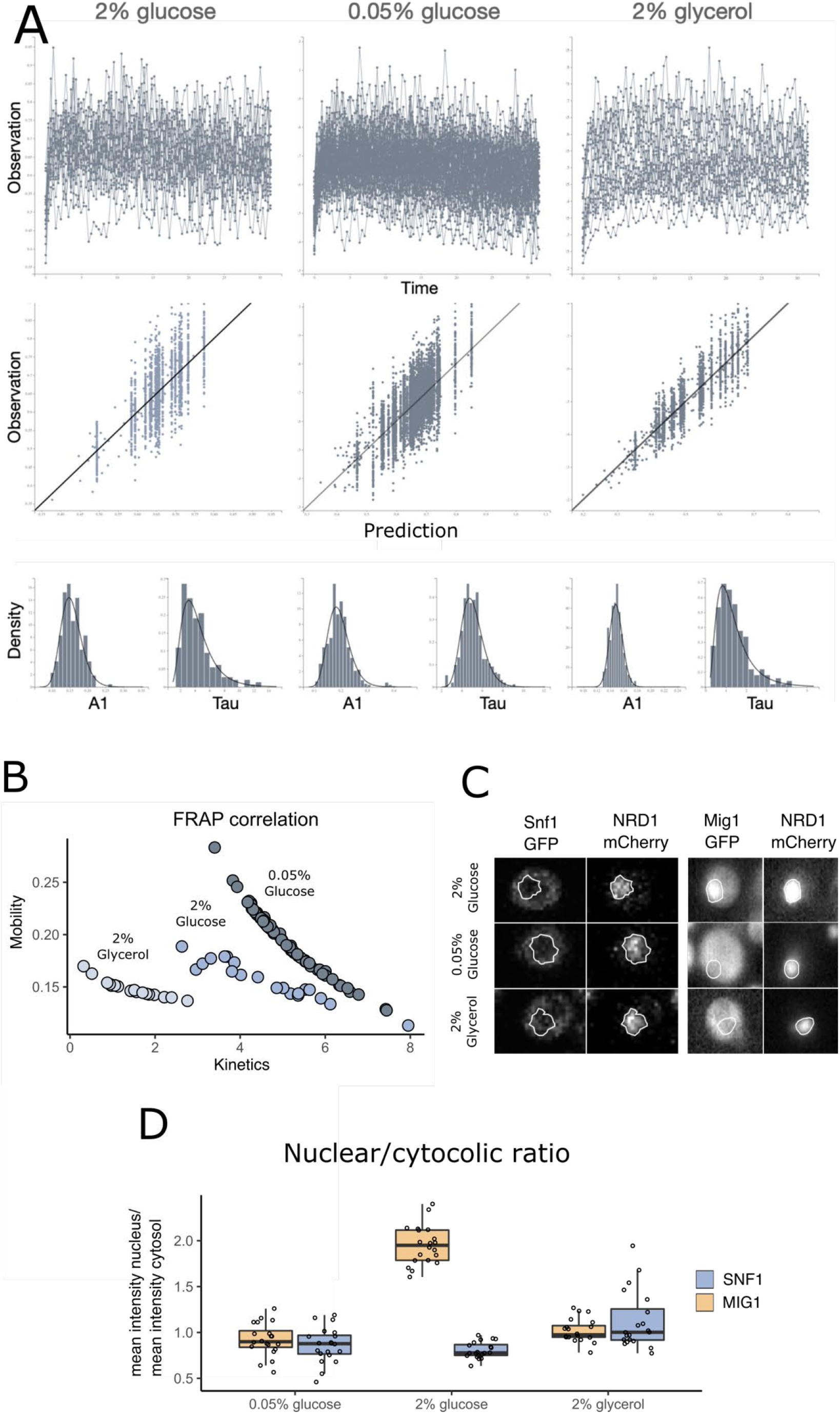
FRAP (Snf1) and nuclear localization (Snf1 and Mig1) measurements of exponentially grown cells exposed to YNB with either 2% glucose, 0.05% glucose or 2% glycerol. (A) the single-cell recovery curves from the FRAP experiment, individual prediction versus observation (IPRED) plot based on the single exponential fit (*I* = *I_0_* + *A1* * (1 – *e^−tau1*t^*), *I_0_* represents the degree of bleaching), and the resulting marginal density plots for both the individual parameters (bar) and population distributions (line) for the parameters *A*1 (the mobility constant) and *tau*1 (the kinetic constant). (B) the correlation between the individual parameters. (C) Snf1-GFP and Mig1-GFP fluorescence relative to the nuclear marker Nrd1-mCherry. The nuclear-to-cytosolic ratio (NC ratio) was calculated by dividing the mean of the fluorescence in the nucleus with the mean of the fluorescence in the cytosol. (D) single-cell nuclear-to-cytoplasmic ratio of Snf1 and Mig1.

Both single and double models were able to describe the data well. However, the Fisher matrix indicates overfitting when using the double exponential equation (summary statistics in SI data files 2, 3 and 4 and complete results at https://github.com/cvijoviclab/Mig1_frap_nlme). The single exponential fit performed best for all conditions (Figure 1A). The 0.05% glucose showed slightly inconsistent behavior with a single exponential curve, but the analysis of the model diagnostics shows that a single-exponential model gives a good approximation of the kinetic behavior.

The fluorescently tagged Snf1 was bleached during the FRAP experiment, and the nuclear fractions differ between conditions, hence the mobility (parameter *A*1) did not accurately reflect the mobile fraction. To correct for the bleached population, we used the steady-state value of the nuclear fraction and the estimated degree of bleaching in each condition (I_0_) to calculate the mobile and immobile fractions of Snf1 (Table 1). The Snf1 steady-state nuclear fractions were similar in high and low glucose (Figure 1D), the mean fold change of 1.06 when comparing 0.05% glucose to 2% glucose, were not significant (p-value = 0.5479). At 2% glycerol the mean NC ratio was increased with a fold change of 1.39 relative 2% glucose (p-value = 1e-05) and 1.30 compared to 0.05% glucose (p-value = 0.0072). Mig1 has a 2.20-fold decrease when comparing the mean NC ratio of 0.05% to 2% glucose (p-value = 4.4e-11) and a 1.98-fold decrease when comparing 2% glycerol to 2% glucose (p-value = 4.4e-11). There was no significant difference of Mig1 nuclear localization between 0.05% glucose and 2% glycerol (p-value = 0.23). Comparing the slight increase in nuclear localization for SNF1 with the effect that the energy availability has for Mig1 nuclear localization it is unlikely that the localization of SNF1 would have a major role in glucose de-repression associated with Mig1. Unlike Snf1, Mig1 nuclear localization is affected by the energy availability, corresponding to SNF1 activity. The slight difference in nuclear localization is also reflected in the kinetic coefficient for Snf1 where the cells grown in glucose show a fast nuclear-cytoplasmic shuttling, in contrast to the cells grown in glycerol showing a slow nuclear-cytoplasmic shuttling (Table 1).

**Table 1:** FRAP population parameters for Snf1 separated by the fixed effects and standard deviation of random effects (W). Correlation between the kinetic constant (tau1) and mobility (A1), the degree of bleaching (I_0_) as well as steady-state data and calculated fractions.

|  | 2% glucose |  | 0.05% glucose |  | 2% glycerol |  |
| --- | --- | --- | --- | --- | --- | --- |
| | $Y_p$ | S.E | $Y_p$ | S.E | $Y_p$ | S.E |
| $A1$ | **0.153** | 0.0177 | **0.191** | 0.00804 | **0.149** | 0.0157 |
| $\tau_{1}$ | **3.93** | 0.961 | **4.98** | 0.18 | **1.18** | 0.221 |
| $I_0$ | **0.501** | 0.0201 | **0.464** | 0.00839 | **0.358** | 0.0245 |
| $W_{A1}$ | **0.184** | 0.108 | **0.209** | 0.0252 | **0.0642** | 0.0464 |
| $W_{\tau_{1}}$ | **0.471** | 0.22 | **0.207** | 0.0255 | **0.592** | 0.139 |
| $W_{I_0}$ | **0.104** | 0.0205 | **0.108** | 0.011 | **0.25** | 0.0406 |
| corr A1-tau1 | **-0.715** | 0.432 | **-0.998** | 0.00175 | **-0.917** | 0.322 |
| $I_{nuc}/I_{cyt}$ | **0.813** | 0.00424 | **0.856** | 0.00494 | **1.117** | 0.00580 |
| Mobile fraction | **0.897** |  | **0.9224** |  | **0.872** |  |
| Immobile fraction | **0.103** |  | **0.0776** |  | **0.128** |  |

The NLME regression approach provides information about cell-to-cell variability and the correlation between population parameters. We observed a strong negative correlation between mobility and the kinetic coefficient (Table 1 and Figure 1B), where cells with a high mobile fraction show a slower kinetic behavior. The opposite relationship was observed between conditions (Figure 1B), where cells grown in 0.05% glucose, with an overall higher mobile fraction, have a general faster kinetic behavior, followed by 2% glucose and 2% glycerol.

### The short-term Mig1 response is more sensitive to glucose than to fructose and mannose

To better understand the sensitivity of the initial Snf1/Mig1 pathway response towards different carbon source levels, cells were exposed to glucose, mannose and fructose. Through fluorescent time-lapse microscopy, the initial spatial response of Mig1 was observed. We exposed cells to concentration shifts from ethanol to 0%, 0.005, 0.05%, 0.5%, 1%, and 4% glucose, mannose or fructose. Mig1 localizing to the nucleus has been observed at 0.005% glucose. While in upshift to mannose and fructose (Figure 3 and Figure S1), Mig1 nuclear localization was only observed at concentrations above 0.05%. These results suggest that the Mig1 nuclear import is more sensitive to glucose than to mannose and fructose. For the upshift to mannose, the maximum nuclear intensity was already reached at the upshift to 0.05% glucose, and the average nuclear intensity did not increase more. Glucose reaches a maximum Mig1 nuclear intensity at a lower concentration compared to fructose. This could be a consequence of the import rates of these hexose sugars since the maximum import rate of glucose is lower than the import rate of fructose (Berthels et al. 2008). Altogether, this suggests that the nuclear import rate of Mig1 is coupled to the import of hexose in the cell or the availability of the hexose sugars inside the cell. Overall, for all hexoses, the Mig1 nuclear intensity increased in a dose-dependent manner, with the rate of increase being hexose specific.

**Figure 3:**
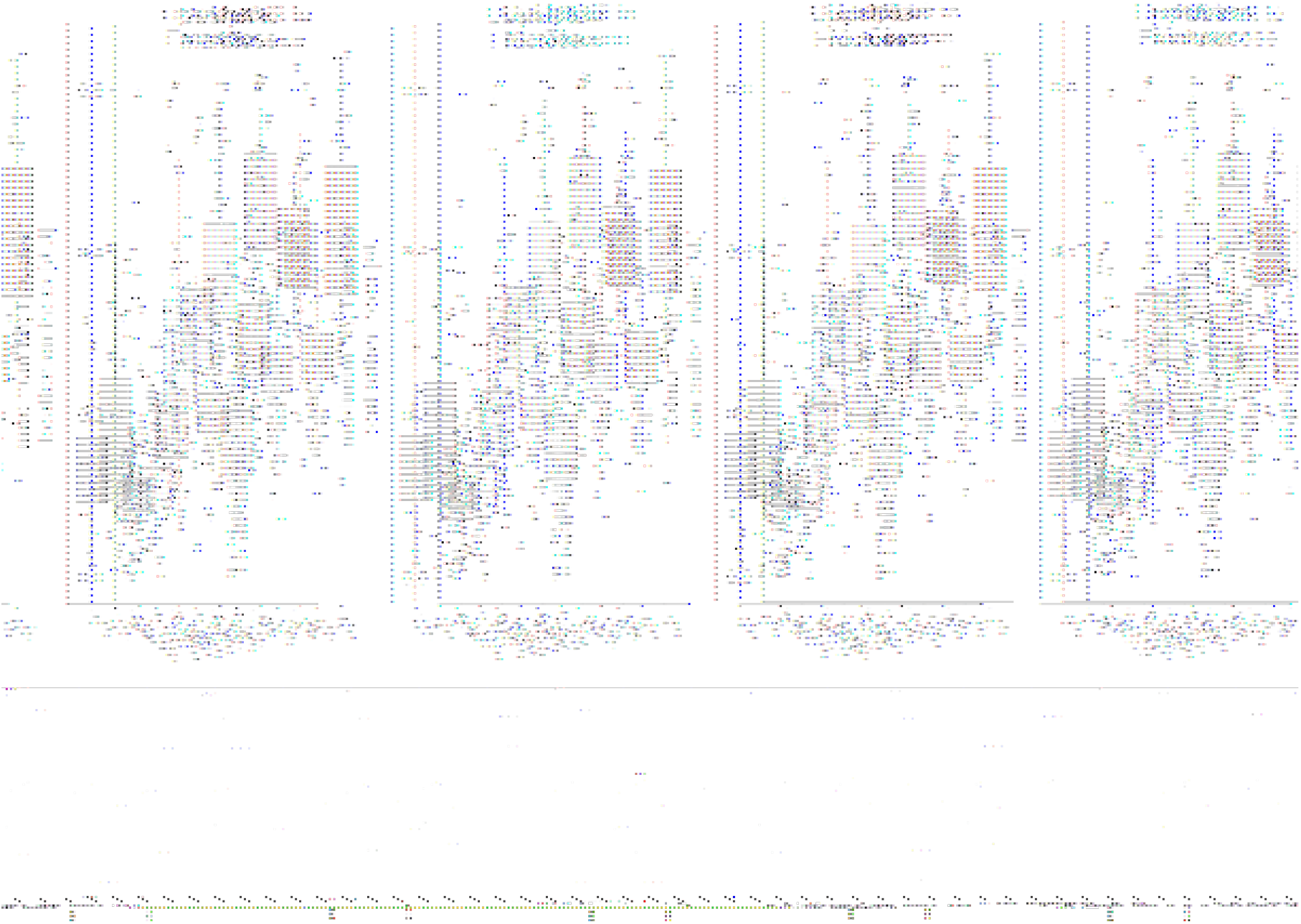
Mig1 nuclear localization in cells after the shift from ethanol to the indicated concentration of glucose, mannose or fructose. Mig1 localization within the observation time of 15 minutes. Each circle represents the maximum Mig1 localization for one cell. Horizontal lines indicate the mean, the boxplot has as lower and upper hinge respectively the 25^th^ and 75^th^ percentile and the whiskers denote the 95% confidence interval. The horizontal dotted gray lines represent either localization index which indicates (bulk) Mig1 located in the nucleus, or cytosol.

## Materials & Methods

### Strains and plasmids

Yeast strains were grown at 30°C in YNB synthetic complete medium containing 1.7 g/l yeast nitrogen base, 5 g/l ammonium sulfate, 670 mg/l complete supplement mix with appropriate drop out where applicable; supplemented with carbon source as indicated by the specific experiments.

#### Strains used in this study

BY4741 *MATa his3Δ1 leu2Δ0 met15Δ0 ura3Δ0*

BY4741 *MATa his3Δ1 leu2Δ0 met15Δ0 ura3Δ0 SNF1-GFP-HIS3 MXNRD1-mCherry-Hph*

W303-1A (202) *MATa {leu2-3,112 trp1-1 can1-100 ura3-1 ade2-1 his3-11,15}*

W303-1A (202) *MATa {leu2-3,112 trp1-1 can1-100 ura3-1 ade2-1 his3-11,15} NRD1-mCherry-Hph MIG1-GFP-KanMX*

### Fluorescent Recovery After Photobleaching (FRAP)

BY4741 and BY4741 *SNF1-GFP-HIS3MX NRD1-mCherry-Hph* were grown in YNB to exponential phase, OD ≈ 0.3, and immobilized on an 8-well Chambered Coverglass (Ibidi) coated with poly-L-lysine (Sigma). Media was switched to YNB supplemented with Complete Supplement Mix (Formedium) with either 2% glucose, 0.05% glucose or 2% glycerol at least 1h before imaging to ensure adaptation to the new carbon source. At least 20 cells per condition were imaged on ELYRA PS.1 SIM/PAL-M LSM780 (Zeiss) using Plan-Apochromat 40x /1.4 oil immersion objective, with settings: 1.59 Airy which equals 1.1 μm z sectioning, 6X zoom with pixel size of 0.28μm and pixel dwell time of 6.14 sec. The cells were continuously imaged for 100 frames, and bleaching was done in 20 bursts at 25% after 10 pre-scans using a circular ROI, of 6 pixels in diameter covering the nucleus.

#### Image processing

The average fluorescent intensity was extracted from the time-lapse image series as well as the time index for each image using the ZEN software (Zeiss). Given the values for background intensity, the intensity for the nuclear region, as well as a control region containing adjacent cells in the same frame, background removal and bleaching correction was done in RStudio (RStudio Team 2020), Version 1.4.1106, and the intensities were normalized based on the pre-scans.

#### Non-linear mixed effect model

A non-linear mixed-effect regression method for analyzing FRAP data was implemented and simulated in Monolix (version 2020R1, Antony, France: Lixoft SAS, 2021 (Kuhn and Lavielle 2005)). The data, project files and models are available at github repository: https://github.com/cvijoviclab/Mig1_frap_n1me.

We used both a double exponential and a single exponential function to fit the data:

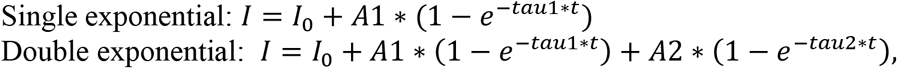

where *I_0_* represent the degree of bleaching, *A*1 is the mobility constant, and *taul* the kinetic constant.

When bleaching the nucleus, a substantial proportion of the fluorescent protein pool is bleached, affecting the calculations of the immobile fraction. To correct for this, the area of the nucleus and the cell was extracted from 20 cells in 0.05% glucose with Fiji software (Schindelin et al. 2012). This was used to calculate cell and nuclear volumes that were assumed to have a spherical shape. The nuclear to cytosolic ratio was calculated based on 20 cells in respective conditions using Fiji for extracting average intensities and R to calculate the ratios by creating a bleaching curve, apply correction and perform linear regression on the time series data. The bleached volume is 2.438379 μm3 and covers the nucleus and assuming Snf1 is mostly vacuole excluded (Chan and Marshall 2014), 8.32% of the cytosol. The immobile fraction derived from the fitted model was then recalculated, accounting for the proportion of the bleached protein pool.

### Steady-state localization microscopy

BY4741 and BY4741 *SNF1-GFP-HIS3MX NRDl-mCherry-Hph* were grown in YNB (Formedium) at 30°C into exponential phase and immobilized on an 8-well Chambered Coverglass (Ibidi) coated with poly-L-lysine (Sigma). Media was switched to YNB with either 2% glucose, 0.05% glucose or 2% glycerol at least 1h before imaging to ensure adaptation to the new carbon source. At least 20 cells/condition were imaged on either ELYRA PS.1 SIM/PAL-M LSM780 (Zeiss) using Plan-Apochromat 40x /1.4 oil immersion objective or DMi8 (Leica) with Lumencor SOLA SE led light (Lumencor) and Leica DFC9000 GT sCMOS camera using HCX PL APO 40x/1.3 oil immersion objective.

#### Image processing

Cell segmentation, extraction of mean intensities and background removal was done in Fiji software and MATLAB _R2019b. Plots and statistical analysis was done using RStudio, Version 1.4.1106.

#### Statistics

As the dataset did not pass the Shapiro–Wilk test, a non-parametric equivalent of ANOVA was used, the Kruskal-Wallis test. For pairwise comparison, a Wilcox test with Bonferroni correction was performed. These statistical tests were done in RStudio, Version 1.4.1106.

### Short-timescale microfluidics experiments

The yeast strains were transformed with GFP-KanMX and mCherry hphNT1 using standard methods for yeast genetics and transformation (Daniel Gietz and Woods 2002). Yeast strains were grown to mid-log phase at 30°C in YNB synthetic complete medium containing 1.7 g/l yeast nitrogen base, 5 g/l ammonium sulfate, 670 mg/l complete supplement mix; 10 mg/l adenine and supplied with 540 mM ethanol overnight. A glass-bottom petri dish (GWST-5030, WillCo Wells, UK) was treated with concanavalin A solution (1 mg/ml in 10mM TrisHCl-buffer, 100mM NaCl, adjusted to pH 8.0 using 5 M HCl) for 30 min at room temperature. The concanavalin A solution was removed, and the cell suspension was added and incubated for 5 min at 30°C. Cells which did not adhere to the surface were removed by washing with YNB. Exposure of cells to different media conditions was performed using a BioPen system (Fluicell AB, Sweden). Experiments were performed on an inverted microscope Olympus cellR widefield microscope system, based on an inverted IX81 motorized microscope with a Xe light source (MT20) and a Hamamatsu C8484 CCD camera. Images were acquired using a U PlanS Apo 40x NA 0.95l objective. The filter cubes, light intensities and exposure time and light intensities for all imaging channels used were as following for GFP: excitation 472/30mm emission 520/35nm with an intensity of 20% for 350 ms. mCherry: excitation 560/40 nm, emission 630/75 nm with an intensity of 20% for 150 ms. The microscope and the microfluidic device were controlled using the Experiment Manager in the Xcellence software. The temperature was set to 30°C. Three images with an axial distance of 0.8 μm were acquired in transmission and fluorescent channels. The acquisition time for one set of images at each time point was ≈15 s. Images were acquired at changing imaging intervals to reduce phototoxicity and bleaching while keeping appropriate timing to monitor changes in Mig1 localization. Time-lapse imaging was performed 3 times every 30 s until the media shift, followed by 15 times every 20 s, followed by 5 times every 120 s, adding up to an overall experiment time of 16 min. Brightfield images acquired above the focal plane were divided by images acquired below the focal plane using custom Matlab scripts. Division of images leads to the elimination of uneven illumination and enhances the diffraction pattern of cells. Segmentation was performed on the resulting images using CellX (C. Mayer et al. 2013). The Mig1-localization index was calculated from the CellX output as follows:

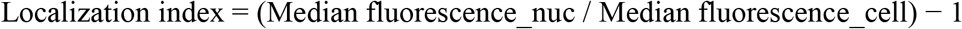

Cells were tracked using custom MATLAB scripts using previously described methods (Ricicova et al. 2013).

## Discussion

In this work, we studied the spatial distribution of Snf1 under different glucose concentrations and the kinetics of the nucleocytoplasmic shuttling by employing a FRAP method. We found that both the spatial distribution of Snf1 and the kinetics of the nucleocytoplasmic shuttling have an equilibrium that show a difference between the type of carbon source, which contrasts with the Snf1 target, Mig1. Using non-linear mixed-effect regression, a negative correlation between Snf1 mobility and the kinetic constant of the nucleocytoplasmic, shuttling was observed within the conditions, while a positive trend was observed between different conditions. This indicates that two different mechanisms are at play, including at least one negative feedback loop. Our time-lapse fluorescent microscopy shows that the intensity of Mig1 localization differs between the same concentration of different hexose sugars in the first 15 minutes after the upshift. Thus, the Snf1/Mig1 pathway seems to be more sensitive to glucose than to fructose and mannose.

Our main observation, that Snf1 show a difference in kinetic behavior and localization based on the presence of glucose and not the activity of Snf1. This is in contrasts with Mig1, which reacts strongly to the glucose concentrations. When we measured the steady-state NC ratio of Snf1 and Mig1 after shifting glucose grown cells to 2% glucose, 0.05% glucose and 2% glycerol, we observed that Mig1 had a large difference in NC ratio between the low and the high glucose concentrations, while the NC ratio in 0.05% glucose and 2% glycerol was similar. The NC ratio of Snf1 does not change significantly between the two glucose concentrations but is significant when comparing the glucose condition to the 2% glycerol, although with a small effect, where we also observe a higher cell-to-cell variability. These data are consistent with previous time-lapse studies on Mig1 where in glucose grown cells shifted to 2% glucose, Mig1 stayed nuclear after 2h. In cells moved to a concentration below 0.2% glucose, Mig1 shifted to the cytosol and remained cytosolic after 2h (Bendrioua et al. 2014). Furthermore, in the parameters fitted from the FRAP recovery curves, the two glucose conditions behave similarly, while the larger difference is between glucose and glycerol. Interestingly, this behavior has been shown in Mig1 nuclear shuttling. The Mig1 kinetics indicate a fast fraction and a slow fraction, where the fast fraction display a short half-time for glucose grown cells and a longer half-time for cells grown in ethanol (Bendrioua et al. 2014), which is interesting since the Mig1 localization is correlated with glucose concentration. However, since both glucose concentrations used where at a level where SNF1 is inactive it cannot be excluded that the difference is corelated with SNF1 activity and not carbon source.

From the parameters inferred by the FRAP recovery curves, a negative correlation between mobility (parameter *A*1) and the kinetic constant (*tau*1) has been observed, indicating the existence of a negative feedback loop. Previous studies suggest the presence of a feedback loop where the concentration of Snf1 inhibit the phosphorylation status of Snf1, and the phosphorylation status is in turn regulating the levels of Snf1 (Hsu et al. 2015). Furthermore, taken in the context where Snf1 needs to be phosphorylated in order to accumulate in the nucleus (Hedbacker, Hong, and Carlson 2004), this can potentially be the negative feedback loop we observe in the FRAP data. One straightforward interpretation is that cells subjected to the same carbon source and concentration but with a large fraction of the Snf1 pool bound to other processes need a higher activity of the nucleocytoplasmic shuttling to serve the same function. Previous studies suggest that the levels and the phosphorylation status of Snf1 are reciprocally regulated, as hyperphosphorylation has been observed when the levels of Snf1 is lower than normal (Hsu et al. 2015). This would fit with a model where nucleocytoplasmic shuttling is regulated by Snf1 phosphorylation status. A lower amount of mobile Snf1 would lead to a higher degree of phosphorylation in the available Snf1 and an increase in nucleocytoplasmic shuttling. As the localization of Snf1 and the pattern of the population kinetic parameters between glucose and glycerol differed, a phosphorylation dependent mechanism would suggest that the cell differentiates between carbon sources upstream SNF1 and act on the rates of phosphorylation and dephosphorylation by the phosphatases Reg1/2-Glc7, Sit4 or Ptc2 and the kinases Elm1, Tos3 and Sak1.

Other mechanisms might also provide explanations as it is not known whether Snf1 mediates its own transport across the nuclear membrane, and it is unclear if Gal83 has a nuclear localization signal (M. C. Schmidt and McCartney 2000). It has been suggested that Snf1 participates in the Mig1 repression complex, also even at low glucose levels. The association of Snf1 to Mig1 repression complex is mediated through Hxk1 or Hxk2 and also contains Mig2, Reg1, Snf4 and Gal83 (Vega et al. 2016). However, this is not a 1:1 ratio as we see a large difference in nuclear intensity, and Mig1 operates in clusters (Wollman et al. 2017). Snf1 might also only participates in a fraction of the complexes formed by the Mig1 clusters, as this only been reported for the SUC2 gene location where both Mig1 and Mig2 co-regulates the gene. Either way, it is possible that Snf1 is co-localizing with other components of this complex and that these components are regulating the nucleoplasm shuttling regarding levels of glucose and type of carbon source. This is supported by the fact that Mig1 localization is affected by *hxk1/2*Δ and that this effect is different depending on both carbon source and level (G. W. Schmidt et al. 2020).

Moreover, we investigated the transient localization of Mig1 after a shift in glucose concentration. For glucose, we observed Mig1 localizing to the nucleus at 0.005% glucose, as reported previously (Bendrioua et al. 2014; Devit, Waddle, and Johnston 1997). For mannose and fructose, we observed Mig1 nuclear localization only at concentrations 0.05% and above. For all hexoses, the Mig1 nuclear intensity increased in a dose-dependent manner, however the rate of increase is hexose specific. This could be caused by different affinity of fructose and mannose to the high affinity hexose transported expressed when grown in a non-fermentable carbon source. HXT7 have been reported to have a twice as high affinity towards glucose than fructose (Reifenberger, Boles, and Ciriacy 1997; Liang and Gaber 1996; Boles and Hollenberg 1997). As the metabolism feedback with Snf1, possible through the hexokinase reaction it is also possible that the rate of metabolizing the different types of sugars also plays a role in the different dynamic behavior of the Mig1 response.

The Snf1/Mig1 pathway is immensely complex, and due to difficulties and lack of experimental methods for monitoring Snf1, the transcription factor Mig1 is often used as a readout. However, previous studies have pointed out that Mig1 is regulated both by a Snf1 dependent and a Snf1 independent mechanism, making it hard to infer the mechanistic behavior of Snf1 by monitoring Mig1. To further elucidate the dynamics and mechanism of the Snf1 pathway, the development of tools for monitoring Snf1, preferably in single cells in yeast, would be needed. For example, a method for monitoring phosphorylation levels without the risk of activating Snf1 or a method to investigate the complexes that Snf1 participates in during different conditions.

In this work, we employed non-linear mixed effect regression to analyze FRAP data, enabling inference of more information than using traditional regression methods. We show that a negative feedback loop controls Snf1 nucleocytoplasmic shuttling. Further, we hypothesize that a mechanism acting upstream Snf1 regulate the rate of phosphorylation/dephosphorylation, thus conferring information on carbon source. This help giving the Snf1-Mig1 system the flexibility and sensitivity to fine-tune itself dynamically to the metabolic state of the cell.

## Acknowledgments

We acknowledge the Cvijovic group members for input and support along the way. We acknowledge the Centre for Cellular Imaging at the University of Gothenburg and National Microscopy Infrastructure, NMI (VR-RFI 2016-00968), for giving us access to high-resolution techniques and excellent guidance in the world of imaging.

## Supplementary Information

**Figure S1.**
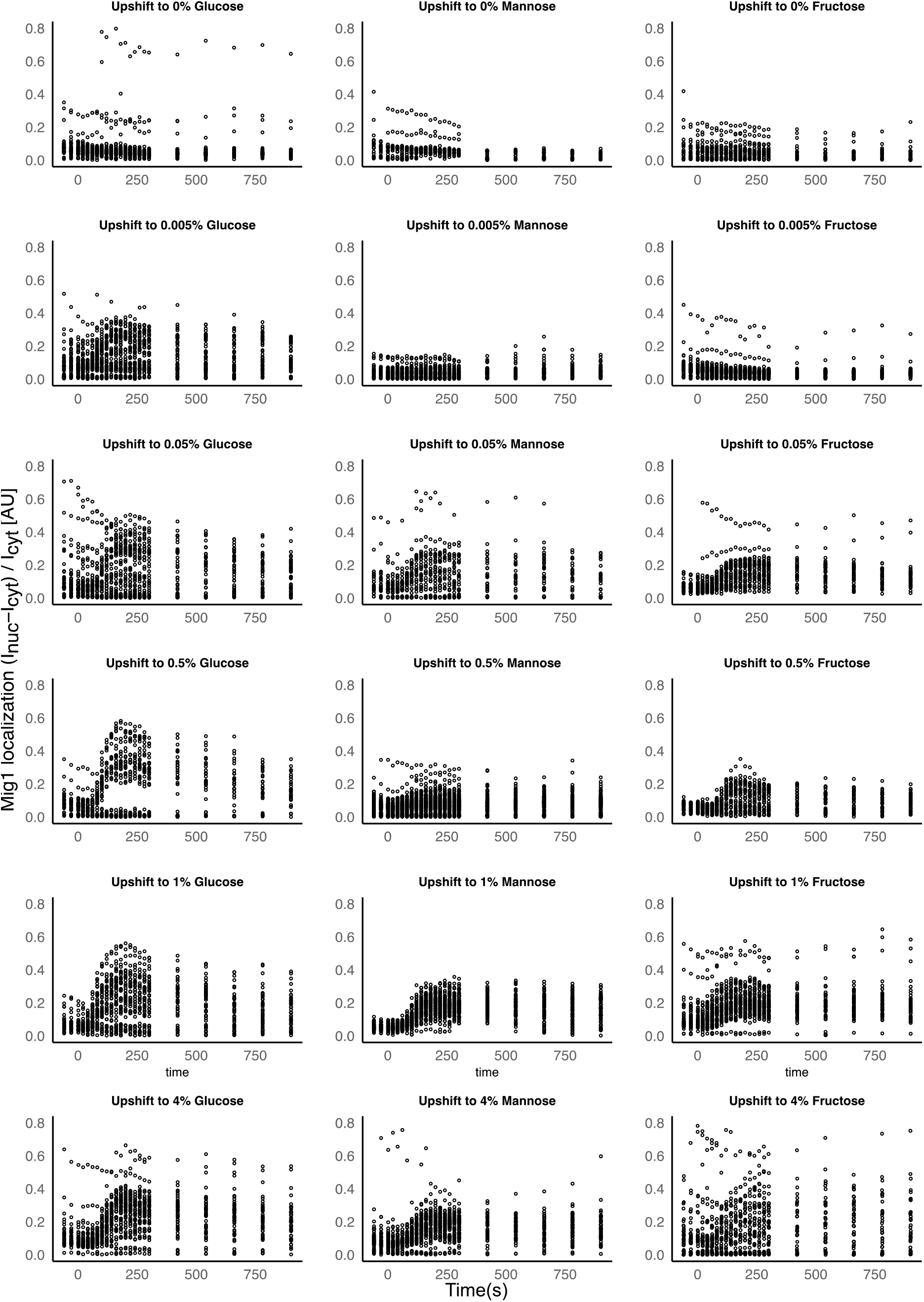
Supplementary data: Mig1 nuclear localization index in WT cells within 15 minutes after shift from ethanol to the indicated concentration of glucose, mannose or fructose (0%, 0.005%, 0.05%, % 0.5%, 1% and 4%).

## References

Almquist, Joachim, Loubna Bendrioua, Caroline Beck Adiels, Mattias Goksör, Stefan Hohmann, and Mats Jirstrand. 2015. “A Nonlinear Mixed Effects Approach for Modeling the Cell-To-Cell Variability of Mig1 Dynamics in Yeast.” PLOS ONE 10 (4): e0124050. https://doi.org/10.1371/journal.pone.0124050.

Bendrioua, Loubna, Maria Smedh, Joachim Almquist, Marija Cvijovic, Mats Jirstrand, Mattias Goksör, Caroline B. Adiels, and Stefan Hohmann. 2014. “Yeast AMP-Activated Protein Kinase Monitors Glucose Concentration Changes and Absolute Glucose Levels.” Journal of Biological Chemistry 289 (18): 12863–75. https://doi.org/10.1074/jbc.m114.547976.

Berthels, Nele J., Ricardo R. Cordero Otero, Florian F. Bauer, Isak S. Pretorius, and Johan M. Thevelein. 2008. “Correlation between Glucose/Fructose Discrepancy and Hexokinase Kinetic Properties in Different Saccharomyces Cerevisiae Wine Yeast Strains.” Applied Microbiology and Biotechnology 77 (5): 1083–91. https://doi.org/10.1007/s00253-007-1231-2.

Boles, Eckhard, and Cornelis P Hollenberg. 1997. “The Molecular Genetics of Hexose Transport in Yeasts.” FEMS Microbiology Reviews 21 (1): 85–111. https://doi.org/10.1111/j.1574-6976.1997.tb00346.x.

Celenza, J L, F J Eng, and M Carlson. 1989. “Molecular Analysis of the SNF4 Gene of Saccharomyces Cerevisiae: Evidence for Physical Association of the SNF4 Protein with the SNF1 Protein Kinase.” Molecular and Cellular Biology 9 (11): 5045–54. https://doi.org/10.1128/mcb.9.11.5045.

Chan, Yee-Hung Mark, and Wallace F. Marshall. 2014. “Organelle Size Scaling of the Budding Yeast Vacuole Is Tuned by Membrane Trafficking Rates.” Biophysical Journal 106 (9): 1986–96. https://doi.org/10.1016/j.bpj.2014.03.014.

Chandrashekarappa, Dakshayini G., Rhonda R. McCartney, Allyson F. O’Donnell, and Martin C. Schmidt. 2016. “The β Subunit of Yeast AMP-Activated Protein Kinase Directs Substrate Specificity in Response to Alkaline Stress.” Cellular Signalling 28 (12): 1881–93. https://doi.org/10.1016/j.cellsig.2016.08.016.

Chandrashekarappa, Dakshayini G., Rhonda R. McCartney, and Martin C. Schmidt. 2013. “Ligand Binding to the AMP-Activated Protein Kinase Active Site Mediates Protection of the Activation Loop from Dephosphorylation*,.” Journal of Biological Chemistry 288 (1): 89–98. https://doi.org/10.1074/jbc.M112.422659.

Daniel Gietz, R., and Robin A. Woods. 2002. “Transformation of Yeast by Lithium Acetate/Single-Stranded Carrier DNA/Polyethylene Glycol Method.” Guide to Yeast Genetics and Molecular and Cell Biology - Part B, 87–96. https://doi.org/10.1016/s0076-6879(02)50957-5.

Davidian, Marie, and David M. Giltinan. 2003. “Nonlinear Models for Repeated Measurement Data: An Overview and Update.” Journal of Agricultural, Biological, and Environmental Statistics 8 (4): 387–419. https://doi.org/10.1198/1085711032697.

DeVit, Michael J., and Mark Johnston. 1999. “The Nuclear Exportin Msn5 Is Required for Nuclear Export of the Mig1 Glucose Repressor of Saccharomyces Cerevisiae.” Current Biology 9 (21): 1231–41. https://doi.org/10.1016/s0960-9822(99)80503-x.

Devit, Michael J., James A. Waddle, and Mark Johnston. 1997. “Regulated Nuclear Translocation of the Mig1 Glucose Repressor.” Molecular Biology of the Cell 8 (8): 1603–18. https://doi.org/10.1091/mbc.8.8.1603.

García-Salcedo, Raúl, Timo Lubitz, Gemma Beltran, Karin Elbing, Ye Tian, Simone Frey, Olaf Wolkenhauer, Marcus Krantz, Edda Klipp, and Stefan Hohmann. 2014. “Glucose De-Repression by Yeast AMP-Activated Protein Kinase SNF1 Is Controlled via at Least Two Independent Steps.” FEBS Journal 281 (7): 1901–17. https://doi.org/10.1111/febs.12753.

Hardie, D. Grahame. 2014. “AMPK—Sensing Energy While Talking to Other Signaling Pathways.” Cell Metabolism 20 (6): 939–52. https://doi.org/10.1016/j.cmet.2014.09.013.

Hardie, D. Grahame, Fiona A. Ross, and Simon A. Hawley. 2012. “AMP-Activated Protein Kinase: A Target for Drugs Both Ancient and Modern.” Chemistry & Biology 19 (10): 1222–36. https://doi.org/10.1016/j.chembiol.2012.08.019.

Hedbacker, Kristina, and Marian Carlson. 2008. “SNF1/AMPK Pathways in Yeast.” Frontiers in Bioscience 13 (13): 2408. https://doi.org/10.2741/2854.

Hedbacker, Kristina, Seung-Pyo Hong, and Marian Carlson. 2004. “Pak1 Protein Kinase Regulates Activation and Nuclear Localization of Snf1-Gal83 Protein Kinase.” Molecular and Cellular Biology 24 (18): 8255–63. https://doi.org/10.1128/mcb.24.18.8255-8263.2004.

Hong, S.-P., F. C. Leiper, A. Woods, D. Carling, and M. Carlson. 2003. “Activation of Yeast Snf1 and Mammalian AMP-Activated Protein Kinase by Upstream Kinases.” Proceedings of the National Academy of Sciences 100 (15): 8839–43. https://doi.org/10.1073/pnas.1533136100.

Hsu, Hsiang En, Tzu Ning Liu, Chung Shu Yeh, Tien Hsien Chang, Yi Chen Lo, and Cheng Fu Kao. 2015. “Feedback Control of Snf1 Protein and Its Phosphorylation Is Necessary for Adaptation to Environmental Stress.” Journal of Biological Chemistry 290 (27): 16786–96. https://doi.org/10.1074/jbc.M115.639443.

Jiang, R, and M Carlson. 1997. “The Snf1 Protein Kinase and Its Activating Subunit, Snf4, Interact with Distinct Domains of the Sip1/Sip2/Gal83 Component in the Kinase Complex.” Molecular and Cellular Biology 17 (4): 2099–2106. https://doi.org/10.1128/mcb.17.4.2099.

Keleher, Cynthia A., Michael J. Redd, Janet Schultz, Marian Carlson, and Alexander D. Johnson. 1992. “Ssn6-Tup1 Is a General Repressor of Transcription in Yeast.” Cell 68 (4): 709–19. https://doi.org/10.1016/0092-8674(92)90146-4.

Kuhn, E., and M. Lavielle. 2005. “Maximum Likelihood Estimation in Nonlinear Mixed Effects Models.” Computational Statistics & Data Analysis 49 (4): 1020–38. https://doi.org/10.1016/j.csda.2004.07.002.

Lavielle, Marc, and France Mentré. 2007. “Estimation of Population Pharmacokinetic Parameters of Saquinavir in HIV Patients with the MONOLIX Software.” Journal of Pharmacokinetics and Pharmacodynamics 34 (2): 229–49. https://doi.org/10.1007/s10928-006-9043-z.

Liang, H, and R F Gaber. 1996. “A Novel Signal Transduction Pathway in Saccharomyces Cerevisiae Defined by Snf3-Regulated Expression of HXT6.” Molecular Biology of the Cell 7 (12): 1953–66.

Llamosi, Artémis, Andres M. Gonzalez-Vargas, Cristian Versari, Eugenio Cinquemani, Giancarlo Ferrari-Trecate, Pascal Hersen, and Gregory Batt. 2016. “What Population Reveals about Individual Cell Identity: Single-Cell Parameter Estimation of Models of Gene Expression in Yeast.” PLOS Computational Biology 12 (2): e1004706. https://doi.org/10.1371/journal.pcbi.1004706.

Mayer, Christian, Sotiris Dimopoulos, Fabian Rudolf, and Jörg Stelling. 2013. “Using CellX to Quantify Intracellular Events.” In Current Protocols in Molecular Biology. John Wiley & Sons, Inc. https://doi.org/10.1002/0471142727.mb1422s101.

Mayer, Faith V., Richard Heath, Elizabeth Underwood, Matthew J. Sanders, David Carmena, Rhonda R. McCartney, Fiona C. Leiper, et al. 2011. “ADP Regulates SNF1, the Saccharomyces Cerevisiae Homolog of AMP-Activated Protein Kinase.” Cell Metabolism 14 (5): 707–14. https://doi.org/10.1016/j.cmet.2011.09.009.

Nath, Nandita, Rhonda R. McCartney, and Martin C. Schmidt. 2003. “Yeast Pak1 Kinase Associates with and Activates Snf1”. Molecular and Cellular Biology 23 (11): 3909–17. https://doi.org/10.1128/mcb.23.11.3909-3917.2003.

Ostling, Jonas, and Hans Ronne. 1998. “Negative Control of the Mig1p Repressor by Snf1p-Dependent Phosphorylation in the Absence of Glucose.” European Journal of Biochemistry 252 (1): 162–68. https://doi.org/10.1046/j.1432-1327.1998.2520162.x.

Persson, Sebastian, Niek Welkenhuysen, Sviatlana Shashkova, and Marija Cvijovic. 2020. “Fine-Tuning of Energy Levels Regulates SUC2 via a SNF1-Dependent Feedback Loop.” Frontiers in Physiology 11. https://doi.org/10.3389/fphys.2020.00954.

Reifenberger, Elke, Eckhard Boles, and Michael Ciriacy. 1997. “Kinetic Characterization of Individual Hexose Transporters of Saccharomyces Cerevisiae and Their Relation to the Triggering Mechanisms of Glucose Repression.” European Journal of Biochemistry 245 (2): 324–33. https://doi.org/10.1111/j.1432-1033.1997.00324.x.

Ricicova, M., M. Hamidi, A. Quiring, A. Niemisto, E. Emberly, and C. L. Hansen. 2013. “Dissecting Genealogy and Cell Cycle as Sources of Cell-to-Cell Variability in MAPK Signaling Using High-Throughput Lineage Tracking.” Proceedings of the National Academy of Sciences 110 (28): 11403–8. https://doi.org/10.1073/pnas.1215850110.

RStudio Team. 2020. RStudio: Integrated Development Environment for R. Boston, MA: RStudio, PBC. http://www.rstudio.com/.

Ruiz, A., X. Xu, and M. Carlson. 2011. “Roles of Two Protein Phosphatases, Reg1-Glc7 and Sit4, and Glycogen Synthesis in Regulation of SNF1 Protein Kinase.” Proceedings of the National Academy of Sciences 108 (16): 6349–54. https://doi.org/10.1073/pnas.1102758108.

Ruiz, Amparo, Xinjing Xu, and Marian Carlson. 2013. “Ptc1 Protein Phosphatase 2C Contributes to Glucose Regulation of SNF1/AMP-Activated Protein Kinase (AMPK) in Saccharomyces Cerevisiae.” Journal of Biological Chemistry 288 (43): 31052–58. https://doi.org/10.1074/jbc.m113.503763.

Schindelin, Johannes, Ignacio Arganda-Carreras, Erwin Frise, Verena Kaynig, Mark Longair, Tobias Pietzsch, Stephan Preibisch, et al. 2012. “Fiji: An Open-Source Platform for Biological-Image Analysis.” Nature Methods 9 (7): 676–82. https://doi.org/10.1038/nmeth.2019.

Schmidt, Gregor W., Niek Welkenhuysen, Tian Ye, Marija Cvijovic, and Stefan Hohmann. 2020. “Mig1 Localization Exhibits Biphasic Behavior Which Is Controlled by Both Metabolic and Regulatory Roles of the Sugar Kinases.” Molecular Genetics and Genomics 295 (6): 1489–1500. https://doi.org/10.1007/s00438-020-01715-4.

Schmidt, M. C., and Rhonda R. McCartney. 2000. “Beta-Subunits of Snf1 Kinase Are Required for Kinase Function and Substrate Definition.” The EMBO Journal 19 (18): 4936–43. https://doi.org/10.1093/emboj/19.18.4936.

Sissoko, Daouda, Cedric Laouenan, Elin Folkesson, Abdoul-Bing M’Lebing, Abdoul-Habib Beavogui, Sylvain Baize, Alseny-Modet Camara, et al. 2016. “Experimental Treatment with Favipiravir for Ebola Virus Disease (the JIKI Trial): A Historically Controlled, Single-Arm Proof-of-Concept Trial in Guinea.” PLOS Medicine 13 (3): e1001967. https://doi.org/10.1371/journal.pmed.1001967.

Treitel, M. A., and M. Carlson. 1995. “Repression by SSN6-TUP1 Is Directed by MIG1, a Repressor/Activator Protein.” Proceedings of the National Academy of Sciences 92 (8): 3132–36. https://doi.org/10.1073/pnas.92.8.3132.

Treitel, Michelle A., Sergei Kuchin, and Marian Carlson. 1998. “Snf1 Protein Kinase Regulates Phosphorylation of the Mig1 Repressor in Saccharomyces Cerevisiae.” Molecular and Cellular Biology 18 (11): 6273–80. https://doi.org/10.1128/mcb.18.11.6273.

Usaite, Renata, Michael C Jewett, Ana Paula Oliveira, John R Yates, Lisbeth Olsson, and Jens Nielsen. 2009. “Reconstruction of the Yeast Snf1 Kinase Regulatory Network Reveals Its Role as a Global Energy Regulator.” Molecular Systems Biology 5 (1): 319. https://doi.org/10.1038/msb.2009.67.

Vega, Montserrat, Alberto Riera, Alejandra Fernández-Cid, Pilar Herrero, and Fernando Moreno. 2016. “Hexokinase 2 Is an Intracellular Glucose Sensor of Yeast Cells That Maintains the Structure and Activity of Mig1 Protein Repressor Complex *.” https://doi.org/10.1074/jbc.M115.711408.

Vincent, O, R Townley, S Kuchin, and M Carlson. 2001. “Subcellular Localization of the Snf1 Kinase Is Regulated by Specific Beta Subunits and a Novel Glucose Signaling Mechanism.” Genes & Development 15 (9): 1104–14. https://doi.org/10.1101/gad.879301.

Welkenhuysen, Niek, Johannes Borgqvist, Mattias Backman, Loubna Bendrioua, Mattias Goksör, Caroline B Adiels, Marija Cvijovic, and Stefan Hohmann. 2017. “Single-Cell Study Links Metabolism with Nutrient Signaling and Reveals Sources of Variability.” BMC Systems Biology 11 (1). https://doi.org/10.1186/s12918-017-0435-z.

Wollman, Adam J.M., Sviatlana Shashkova, Erik G. Hedlund, Rosmarie Friemann, Stefan Hohmann, and Mark C. Leake. 2017. “Transcription Factor Clusters Regulate Genes in Eukaryotic Cells.” ELife 6 (August). https://doi.org/10.7554/eLife.27451.

Zhang, Jie, Lisbeth Olsson, and Jens Nielsen. 2010. “The β-Subunits of the Snf1 Kinase in Saccharomyces Cerevisiae, Gal83 and Sip2, but Not Sip1, Are Redundant in Glucose Derepression and Regulation of Sterol Biosynthesis.” Molecular Microbiology 77 (2): 371–83. https://doi.org/10.1111/j.1365-2958.2010.07209.x.

Zhang, Yuxun, Rhonda R. McCartney, Dakshayini G. Chandrashekarappa, Simmanjeet Mangat, and Martin C. Schmidt. 2011. “Reg1 Protein Regulates Phosphorylation of All Three Snf1 Isoforms but Preferentially Associates with the Gal83 Isoform.” Eukaryotic Cell 10 (12): 1628–36. https://doi.org/10.1128/EC.05176-11.

