## Supplementary files for "Localization and phosphorylation in the Snf1 network is controlled by two independent pathways"

```
********************************************************************************
*                    Double_exp_mac.mlxtran                    *
*                    May 18, 2021 at 10:53:18                  *
*                    Monolix version : 2020R1                  *
********************************************************************************
```

### ESTIMATION OF THE POPULATION PARAMETERS ________________________________________

| Fixed Effects --------------------------- |  |  | se_sa | rse(%) |
|---|---|---|---|---|
| A1_pop | : | 0.0498 | nan | nan |
| tau1_pop | : | 1.98e+3 | nan | nan |
| A2_pop | : | 0.114 | nan | nan |
| tau2_pop | : | 3.39 | nan | nan |
| I0_pop | : | 0.449 | 0.00966 | 2.15 |

| Standard Deviation of the Random Effects - | | | | |
|---|---|---|---|---|
| omega_A1 | : | 0.381 | 0.0618 | 16.2 |
| omega_tau1 | : | 3.66 | 0.317 | 8.67 |
| omega_A2 | : | 0.231 | 0.0285 | 12.3 |
| omega_tau2 | : | 1.27 | 0.249 | 19.7 |
| omega_I0 | : | 0.194 | 0.0156 | 8.03 |

| Correlations ---------------------------- |  |  |  |  |
|---|---|---|---|---|
| corr_A2_A1 | : | 0.973 | 0.0112 | 1.15 |
| corr_tau1_A1 | : | -0.912 | 0.0449 | 4.92 |
| corr_tau2_A1 | : | -0.946 | 0.0248 | 2.62 |
| corr_tau1_A2 | : | -0.98 | 0.0142 | 1.45 |
| corr_tau2_A2 | : | -0.853 | 0.0577 | 6.76 |
| corr_tau2_tau1 | : | 0.758 | 0.0952 | 12.6 |

| Error Model Parameters ------------------- |  |  |  |  |
|---|---|---|---|---|
| a | : | 0.0208 | 0.00355 | 17.1 |
| b | : | 0.0667 | 0.00568 | 8.52 |

Elapsed time (seconds) : 53
CPU time (seconds) : 94

________________________________________________________________________________

### ESTIMATION OF THE INDIVIDUAL PARAMETERS ________________________________________

Estimation of the individual parameters by Conditional Distribution ------------

| | | min | Q1 | median | Q3 | max |
|---|---|---|---|---|---|---|
| A1 | : | 0.0167 | 0.0423 | 0.0509 | 0.0598 | 0.108 |
| tau1 | : | 2.86 | 5.28e+3 | 2.73e+4 | 1.82e+5 | 1.42e+7 |
| A2 | : | 0.0607 | 0.104 | 0.115 | 0.128 | 0.182 |
| tau2 | : | 0.152 | 2.61 | 4.66 | 8.59 | 122 |
| I0 | : | 0.257 | 0.412 | 0.473 | 0.506 | 0.613 |

Elapsed time (seconds) : 16
CPU time (seconds) : 34

-----------------------------------------------------------------------------

Estimation of the individual parameters by Conditional Mode --------------------

| | min | Q1 | median | Q3 | max |
|---|---|---|---|---|---|
| A1 : | 0.0195 | 0.0434 | 0.0565 | 0.0646 | 0.129 |
| tau1 : | 1.4 | 178 | 696 | 8.36e+3 | 1.27e+7 |
| A2 : | 0.0644 | 0.104 | 0.123 | 0.135 | 0.178 |
| tau2 : | 0.0637 | 1.51 | 2.26 | 4.89 | 55.6 |
| I0 : | 0.254 | 0.405 | 0.463 | 0.5 | 0.613 |

Elapsed time (seconds) : 1.7
CPU time (seconds) : 5

-----------------------------------------------------------------------------

______________________________________________________________________________

ESTIMATION OF THE FISHER INFORMATION MATRIX ___________________________________

Estimation of the Fisher information matrix by Stochastic Approximation --------

Correlation Matrix :
A1_pop nan
tau1_pop nan nan
A2_pop nan nan nan
tau2_pop nan nan nan nan
I0_pop nan nan nan nan 1
omega_A1 nan nan nan nan -0.045251 1
corr_tau1_A1 nan nan nan nan 0.053178 -0.8371 1
corr_A2_A1 nan nan nan nan -0.044081 0.79065 -0.99191 1
corr_tau2_A1 nan nan nan nan 0.061913 -0.89384 0.97509 -0.95076 1
omega_tau1 nan nan nan nan 0.0075125 0.83379 -0.57028 0.5539 -0.59743 1
corr_tau1_A2 nan nan nan nan 0.065851 -0.89086 0.98023 -0.948 0.98901 -0.58568 1
corr_tau2_tau1 nan nan nan nan -0.042597 0.786 -0.98961 0.99933 -0.94945 0.55459 -0.94348
1
omega_A2 nan nan nan nan -0.02988 0.98362 -0.78629 0.74852 -0.83255 0.91882 -0.82821
0.7448 1
corr_tau2_A2 nan nan nan nan 0.049638 -0.83413 0.99414 -0.9928 0.97933 -0.57824 0.96852
-0.99294 -0.78568 1
omega_tau2 nan nan nan nan -0.052774 0.99392 -0.84174 0.78825 -0.90693 0.77586 -0.90549
0.78384 0.95943 -0.83679 1
omega_I0 nan nan nan nan -0.011701 -0.042447 0.0285 -0.021705 0.035812 -0.015385 0.038481
-0.020201 -0.035316 0.025837 -0.04634 1
a nan nan nan nan-0.0027822 -0.024398 0.011454-0.0084093 0.01679 -0.024647 0.016322-
0.0083925 -0.024873 0.011558 -0.025088 0.0042616 1
b nan nan nan nan 0.0025777 0.01676-0.0042529 0.0016625-0.0090217 0.019628-0.0085718
0.001662 0.0178 -0.004377 0.01729-0.0040679 -0.98865 1

WARNING : Impossible to compute the eigen values of the correlation matrix.

Elapsed time (seconds) : 4.6
CPU time (seconds) : 11

---

ESTIMATION OF THE LOG-LIKELIHOOD

| | | |
|---|---|---|
| | | (is) |
| -2 x log-likelihood | (OFV) : | -18132.51 |
| Akaike Information Criteria | (AIC) : | -18096.51 |
| Corrected Bayesian Information Criteria | (BICc) : | -18021.24 |
| Bayesian Information Criteria | (BIC) : | -18051.92 |

Elapsed time (seconds) : 32.49
CPU time (seconds) : 66.00

[Importance Sampling] Standard error : 1270.754
Sampling distribution : T-distribution with 5 degrees of freedom

DATASET INFORMATION
Number of individuals: 88
Number of observations (observation): 7040
Number of doses: 0

```
*******************************************************************************
*                    Double_exp_mac.mlxtran                    *
*                    May 18, 2021 at 11:00:54                   *
*                    Monolix version : 2020R1                   *
*******************************************************************************
```

### ESTIMATION OF THE POPULATION PARAMETERS

```
Fixed Effects ----------------------------     se_sa    rse(%)
A1_pop         :                   0.11       nan       nan
tau1_pop       :                   4.04       nan       nan
A2_pop         :                 0.0469       nan       nan
tau2_pop       :                   2.92       nan       nan
I0_pop         :                  0.495    0.0127      2.57

Standard Deviation of the Random Effects -
omega_A1       :                  0.281       nan       nan
omega_tau1     :                  0.356    0.0218      6.12
omega_A2       :                  0.136   0.00974      7.18
omega_tau2     :                  0.574       nan       nan
omega_I0       :                 0.0996    0.0216      21.6

Correlations ----------------------------
corr_A2_A1     :                  0.413    0.0403      9.75
corr_tau1_A1   :                  -0.29    0.0766      26.5
corr_tau2_A1   :                 -0.843    0.0431      5.12
corr_tau1_A2   :                  0.749    0.0398      5.31
corr_tau2_A2   :                -0.0439       nan       nan
corr_tau2_tau1 :                  0.594       nan       nan

Error Model Parameters -------------------
a              :               0.000762    0.0112   1.47e+3
b              :                 0.0947    0.0174      18.4

Elapsed time (seconds) :     36
CPU time     (seconds) :     64
```

---

### ESTIMATION OF THE INDIVIDUAL PARAMETERS

```
Estimation of the individual parameters by Conditional Distribution ------------
          min       Q1    median       Q3      max
A1   :  0.0649   0.0966    0.107    0.128    0.178
tau1 :    3.06     3.88     4.37     4.89     5.64
A2   :  0.0412   0.0458   0.0478   0.0499   0.0537
tau2 :    1.34     2.48     3.59     4.14     9.57
I0   :   0.383     0.48    0.498    0.515    0.576
```

Elapsed time (seconds) : 7.2
CPU time (seconds) : 15

-----------------------------------------------------------------------------

Estimation of the individual parameters by Conditional Mode --------------------

| | min | Q1 | median | Q3 | max |
|---|---|---|---|---|---|
| A1 : | 0.0712 | 0.0986 | 0.11 | 0.133 | 0.174 |
| tau1 : | 2.78 | 3.76 | 3.96 | 4.29 | 4.86 |
| A2 : | 0.0421 | 0.0455 | 0.0471 | 0.0487 | 0.0525 |
| tau2 : | 1.21 | 1.95 | 2.96 | 3.55 | 6.26 |
| I0 : | 0.378 | 0.478 | 0.494 | 0.515 | 0.581 |

Elapsed time (seconds) : 0.33
CPU time (seconds) : 1

-----------------------------------------------------------------------------

______________________________________________________________________________

ESTIMATION OF THE FISHER INFORMATION MATRIX ________________________________

Estimation of the Fisher information matrix by Stochastic Approximation --------

Correlation Matrix :

```
      A1_pop      nan
    tau1_pop      nan      nan
      A2_pop      nan      nan      nan
    tau2_pop      nan      nan      nan      nan
      I0_pop      nan      nan      nan      nan        1
    omega_A1      nan      nan      nan      nan      nan      nan
 corr_tau1_A1      nan      nan      nan      nan -0.096054      nan        1
   corr_A2_A1      nan      nan      nan      nan  0.013641      nan  0.45851        1
 corr_tau2_A1      nan      nan      nan      nan  0.036753      nan  -0.11304   0.5205        1
   omega_tau1      nan      nan      nan      nan  -0.17344      nan  0.30991  -0.37655  -0.67455        1
 corr_tau1_A2      nan      nan      nan      nan  -0.11367      nan   0.8253  -0.11813  -0.51506  0.61018        1
corr_tau2_tau1      nan      nan      nan      nan      nan      nan      nan      nan      nan      nan      nan      nan
    omega_A2      nan      nan      nan      nan  -0.18633      nan  0.65284  0.019826  -0.51503   0.89745   0.74014
nan        1
 corr_tau2_A2      nan      nan      nan      nan      nan      nan      nan      nan      nan      nan      nan      nan      nan
nan
   omega_tau2      nan      nan      nan      nan      nan      nan      nan      nan      nan      nan      nan      nan      nan
nan      nan
    omega_I0      nan      nan      nan      nan  0.01675      nan  0.21872-0.0057416 -0.001878  -0.16416    0.2365
nan  -0.10467      nan      nan        1
           a      nan      nan      nan      nan -0.026125      nan  0.077499 0.0042535 -0.022062  0.093891   0.08256
nan   0.11156      nan      nan -0.025206        1
           b      nan      nan      nan      nan  0.030421      nan   -0.1222-0.00012145  0.034021  -0.11286  -0.13375
nan   -0.1393      nan      nan 0.0058086  -0.99409        1
```

WARNING : Impossible to compute the eigen values of the correlation matrix.

Elapsed time (seconds) : 4.7
CPU time (seconds) : 11

----------------------------------------------------------------------------

---

ESTIMATION OF THE LOG-LIKELIHOOD

| | | |
|---|---|---|
| | (is) | |
| -2 x log-likelihood | (OFV) : | -4761.68 |
| Akaike Information Criteria | (AIC) : | -4725.68 |
| Corrected Bayesian Information Criteria | (BICc) : | -4676.26 |
| Bayesian Information Criteria | (BIC) : | -4707.76 |

| | | |
|---|---|---|
| Elapsed time | (seconds) : | 8.25 |
| CPU time | (seconds) : | 16.00 |

[Importance Sampling] Standard error : 13.246
Sampling distribution : T-distribution with 5 degrees of freedom

---

DATASET INFORMATION
Number of individuals: 20
Number of observations (observation): 1800
Number of doses: 0

```
********************************************************************************
*                  Double_exp_mac.mlxtran                          *
*                  May 18, 2021 at 10:57:43                        *
*                  Monolix version : 2020R1                        *
********************************************************************************

ESTIMATION OF THE POPULATION PARAMETERS ________________________________________

Fixed Effects ----------------------------     se_sa    rse(%)
A1_pop       :                    0.118   0.00667      5.63
tau1_pop     :                     1.22     0.303      24.7
A2_pop       :                   0.0387   0.00532      13.8
tau2_pop     :                     2.82      1.17      41.6
I0_pop       :                    0.346    0.0205      5.94

Standard Deviation of the Random Effects -
omega_A1     :                    0.195    0.0308      15.8
omega_tau1   :                    0.651     0.101      15.5
omega_A2     :                    0.291    0.0492      16.9
omega_tau2   :                    0.906     0.144      15.9
omega_I0     :                    0.255    0.0406      15.9

Correlations ----------------------------
corr_A2_A1     :                  -0.11       nan       nan
corr_tau1_A1   :                 -0.754       nan       nan
corr_tau2_A1   :                  0.233       nan       nan
corr_tau1_A2   :                  0.713       nan       nan
corr_tau2_A2   :                 -0.957       nan       nan
corr_tau2_tau1 :                 -0.815       nan       nan

Error Model Parameters -------------------
a            :                 2.22e-16   0.00458  2.06e+15
b            :                   0.0889    0.0091      10.2

Elapsed time (seconds) :      37
CPU time     (seconds) :      63

________________________________________________________________________________

ESTIMATION OF THE INDIVIDUAL PARAMETERS ________________________________________

Estimation of the individual parameters by Conditional Distribution ------------
           min       Q1    median      Q3       max
A1   :  0.0825    0.107    0.117    0.134    0.189
tau1 :   0.285    0.972     1.57     1.92     2.39
A2   :  0.0235   0.0357   0.0411   0.0445   0.0462
tau2 :    1.91     2.37     3.16     5.72       14
I0   :   0.212    0.274    0.333    0.454    0.506
```

```
Elapsed time (seconds) :     6.2
CPU time     (seconds) :      13

 --------------------------------------------------------------------------

Estimation of the individual parameters by Conditional Mode --------------------
           min       Q1    median       Q3       max
A1   :   0.0969    0.114     0.12    0.137    0.178
tau1 :    0.322    0.731     1.19     1.55     1.85
A2   :   0.0249   0.0348   0.0394   0.0413    0.045
tau2 :     1.77     2.28      2.8     4.57     11.3
I0   :    0.204    0.277     0.33    0.443    0.503

Elapsed time (seconds) :    0.35
CPU time     (seconds) :       1

 --------------------------------------------------------------------------

 ____________________________________________________________________________

ESTIMATION OF THE FISHER INFORMATION MATRIX __________________________________

Estimation of the Fisher information matrix by Stochastic Approximation --------

Correlation Matrix :
      A1_pop        1
    tau1_pop  -0.59443        1
      A2_pop  -0.19346   0.83612        1
    tau2_pop   0.16633  -0.89165  -0.91254        1
      I0_pop -0.062262  -0.12924  -0.19041   0.19356        1
    omega_A1  -0.23502   0.49002   0.45678  -0.46566 -0.086062        1
 corr_tau1_A1      nan      nan      nan      nan      nan      nan      nan
   corr_A2_A1      nan      nan      nan      nan      nan      nan      nan      nan
 corr_tau2_A1      nan      nan      nan      nan      nan      nan      nan      nan      nan
   omega_tau1   -0.2332   0.48151   0.44942  -0.45647 -0.083672   0.99909      nan      nan      nan        1
 corr_tau1_A2      nan      nan      nan      nan      nan      nan      nan      nan      nan      nan      nan
corr_tau2_tau1      nan      nan      nan      nan      nan      nan      nan      nan      nan      nan      nan      nan
    omega_A2  -0.17605   0.30921   0.21817  -0.27898 -0.035599   0.65164      nan      nan      nan   0.65282      nan
nan        1
 corr_tau2_A2      nan      nan      nan      nan      nan      nan      nan      nan      nan      nan      nan      nan      nan
nan
   omega_tau2  -0.23148   0.49486   0.46507  -0.47376 -0.088671   0.99873      nan      nan      nan   0.99882      nan
nan   0.63729      nan        1
    omega_I0 0.0066676   0.02698   0.03555 -0.037799 -0.012647 0.0090292      nan      nan      nan  0.010716      nan
nan 0.0055747      nan   0.01065        1
           a 0.0061306 -0.094652  -0.12671   0.11177  0.036509 -0.051481      nan      nan      nan -0.048352      nan
nan -0.018554      nan -0.0512340.00017034        1
           b-0.0069991  0.094798   0.12443  -0.11145 -0.036579  0.056428      nan      nan      nan  0.053322      nan
nan  0.019733      nan  0.056426-0.0012237  -0.98597        1
```

WARNING : Impossible to compute the eigen values of the correlation matrix.

Elapsed time (seconds) : 5.8
CPU time (seconds) : 13

---------------------------------------------------------------------------

---

ESTIMATION OF THE LOG-LIKELIHOOD

| | | |
|---|---|---|
| | | (is) |
| -2 x log-likelihood | (OFV) : | -5905.37 |
| Akaike Information Criteria | (AIC) : | -5869.37 |
| Corrected Bayesian Information Criteria | (BICc) : | -5819.95 |
| Bayesian Information Criteria | (BIC) : | -5851.44 |

| | | |
|---|---|---|
| Elapsed time | (seconds) : | 8.21 |
| CPU time | (seconds) : | 16.00 |

[Importance Sampling] Standard error : 47.242
Sampling distribution : T-distribution with 5 degrees of freedom

---

DATASET INFORMATION
Number of individuals: 20
Number of observations (observation): 1800
Number of doses: 0
